# Folate deficiency facilitates coordination of KDM6A with p53 in response to DNA damage

**DOI:** 10.1101/591768

**Authors:** Jun Gao, Jizhen Zou, Jianting Li, Yongfeng Zhang, Lina Qiao, Fang Wang, Yihua Bao, Jin Hu, Ting Zhang, Qiu Xie, Changyun Liu

## Abstract

Folate contributes to the accumulation of DNA strand breaks (DSBs) in the genome. Kdm6a plays a critical role in early embryogenesis but it is unknown whether Kdm6a is involved in the DNA damage response under folate deficiency. Here, we established a low folate environment for mouse embryonic cells using a folate antagonist (methotrexate, MTX). We found increased enrichment of DSBs in Kdm6a in MTX-treated cells, resulting in reduced Kdm6a expression. MTX treatment enriched KDM6A in the nonhomologous end-joining (NHEJ) repair pathway and controlled the expression of the Ku heterodimer (Ku70/80) and XRCC4. The activation of NHEJ repair pathway-associated genes under folate deficiency relied on the specific interaction between KDM6A and p53. p53 silencing increased Ku heterodimer expression. In addition, in a neural tube defect (NTD) mouse model and low folate neural tube defect human brain samples, KDM6A levels were also decreased and accompanied by over-expression of Ku80. Our findings highlight how alterations in folate levels affect KDM6A with respect to DNA breakage and DNA repair, offering a new insight into the molecular function of KDM6A.

**Summary statement:** Our study demonstrates for the first time that in mammals, Kdm6a and p53 have synergistic effects during DNA fragmentation repair.

## Introduction

Neural tube defects (NTDs) are a common birth defect that can cause infant mortality and severe birth defects. NTDs result from the failure of neural tube to close during early development. The etiology of NTDs is associated with both genetic and environmental factors (Wallingford et al., 2013). The most established environmental factor is folate; pre-pregnancy supplementation with folate can reduce the occurrence of NTDs by 30%–70% (Czeizel and Dudas, 1992). However, the mechanism for this action remains largely unknown. Accumulating evidence shows that folate plays a key role in ensuring maintenance of genomic integrity and normal cell proliferation, especially during early embryonic development (Qiu et al., 2017).

DNA double strand breaks (DSBs) are one of the most deleterious forms of damage that a genome can suffer. Genetic and environmental factors both contribute to structural damage to DNA, and if this damage is not repaired in time, gene expression may be affected, leading to abnormal development. Single-nucleotide polymorphisms (SNPs) in key DNA repair genes may increase the risk of spina bifida, such as *Xeroderma pigmentosum D (XPD), DNA ligase III (LIG3) (APE1)*, and *X-ray repair cross-compleme*nting *3 (XRCC3*). Mutation of DNA-dependent protein kinase catalytic subunit (DNA-PKcs) results in embryonic lethality associated with neuronal apoptosis (Aravinthan et al. 2012; Wang et al. 2008; Li et al. 2018). Folate is a key determinant of DNA strand breaks because of uracil misincorporation, leading to genetic instability (Blount et al, 1997). Also, there is significant DNA damage in tissues of NTD cases and the amount of damage increases with severity of the disease (Aravinthan et al. 2012). DSBs can be repaired by two pathways in mammalian cells: homologous recombination (HR) and nonhomologous end-joining (NHEJ). The former accurately repairs DSBs through homologous DNA sequences and is restricted to S/G2 phases of the cell cycle, while the latter directly ligates DSBs without homology and functions in all phases of the cell cycle. NHEJ is the primary DNA repair pathway for DSBs in multicellular eukaryotes. The core machinery for NHEJ is typically composed of a Ku heterodimer (Ku80/70, XRCC5/6), DNA-PKcs, DNA Ligase IV, XRCC4, and the XRCC4-like factor (XLF). The Ku70/80 heterodimer initially recognizes and binds to DSBs and then DNA-PKcs is recruited to form an active DNA-PK complex (Wang et al. 2008). Finally, XRCC4/DNA Ligase IV performs the repair.

Lysine Demethylase 6A (KDM6A) contains a JmjC-domain. KDM6A is a histone demethylase that removes a methyl group from H3K27. *Kdm6a* knockout (KO) studies have unraveled important roles in many developmental processes, including animal body patterning by regulation of homeobox (HOX) genes (Agger et al., 2007; Lee et al., 2007; Van et al., 2014). Defective anteroposterior patterning may result in NTDs. KDM6A is a general factor that regulates gene transcription during development. However, an additional essential role of KDM6A has been found in the DNA damage response in *Drosophila*. Demethylation by KDM6A regulates Ku80 expression in a p53-dependent manner after exposure to ionizing radiation (IR) (Zhang et al., 2013). Folate can cause epigenetic modification of methylation modifications, including histone methylation (Tibbetts et al., 2010; Martin and Zhang, 2005; Park et al., 2012). However, whether KDM6A is involved in DNA strand breaks and genetic instability in folate-deficient conditions during early development is not known. Therefore, we explored whether KDM6A is involved in the regulation of NHEJ repair in mouse embryonic stem cells (mESCs) using a folate antagonist (methotrexate, MTX). We have previously shown that MTX can cause DNA breakage, which indicates it to be a risk factor for NTDs (Wang et al., 2014).

In this study, we first detected a strong and significant enrichment of DSBs in *Kdm6a* in mESCs treated with MTX, resulting in decreased expression of *Kdm6a*. Decreased KDM6A was enriched in NHEJ repair pathway and played an essential role in upregulation of the Ku heterodimer (Ku70/80) and XRCC4 upon MTX treatment. The activation of NHEJ repair pathway-associated genes depended on the specific interaction between KDM6A and p53 under folate-deficient conditions. p53 silencing lead to the reduced Ku heterodimer expression. In addition, in NTD mouse models and low-folate NTD human brain samples, KDM6A was also decreased, and was accompanied by over-expression of Ku80. Our findings highlight how alterations in folate levels affect KDM6A and DNA breakage leading to a DNA repair response, and offer new insight into the molecular function of KDM6A.

## Results

### Folate deficiency leads to decreased expression of *Kdm6a* in mESCs

The *Kdm6a* gene is located in cytogenetic band 11.3 on the short arm of the X chromosome (Fig 1A) and plays an important role in early embryonic development and DNA repair. It has been extensively studied as a specific demethylase of H3K27me3. To explore the role of *Kdm6a* in repairing DNA breaks induced by folate deficiency, we established a folate deficient mESC model. SV129 cells were treated with MTX (0.12 μM) for 24 h, and compared with non-treated cells. After MTX treatment for 24 h the folate concentration in cells was 10.19 ± 0.79ng/10^6^, which was significantly lower than that of untreated cells at 17.35 ± 2.34ng/10^6^ (Fig 1B). This showed that MTX can induce a low folate environment. Then we wanted to assess the presence of DNA breaks. We extracted proteins from the two groups of cells and found that the level of gH2AX, a DNA break marker protein, in the MTX group was double that in the control group by western blot analysis (Fig 1C), indicating that MTX induced DNA breaks. We also performed immunofluorescence staining for gH2AX and found that the fluorescence signal of gH2AX increased after MTX treatment (Fig 1D). We have previously shown that MTX treatment causes DNA breaks at high frequency in the *Kdm6a* gene (Fig 1E). We, therefore, performed chromatin immunoprecipitation (ChIP) for gH2AX and RT-PCR and demonstrated obvious breaks in the *Kdm6a* gene (Fig 1F). To confirm that *Kdm6a* undergoes high-frequency breaks after MTX treatment we explored the level of *Kdm6a* transcription. RT-PCR showed that MTX treatment decreased *Kdm6a* transcription (Fig 1G). We also assessed KDM6A protein levels. Western blot analysis of KDM6A revealed an approximate 25% decrease in the MTX-treated group. (Fig 1H). Immunofluorescence analysis showed a weakened KDM6A signal, indicating reduced expression (Fig 1I). Therefore, we conclude that in mESCs, a low-folate environment induced by MTX causes high-frequency cleavage of the *Kdm6a* gene, resulting in a decrease in expression.

**Fig 1.**
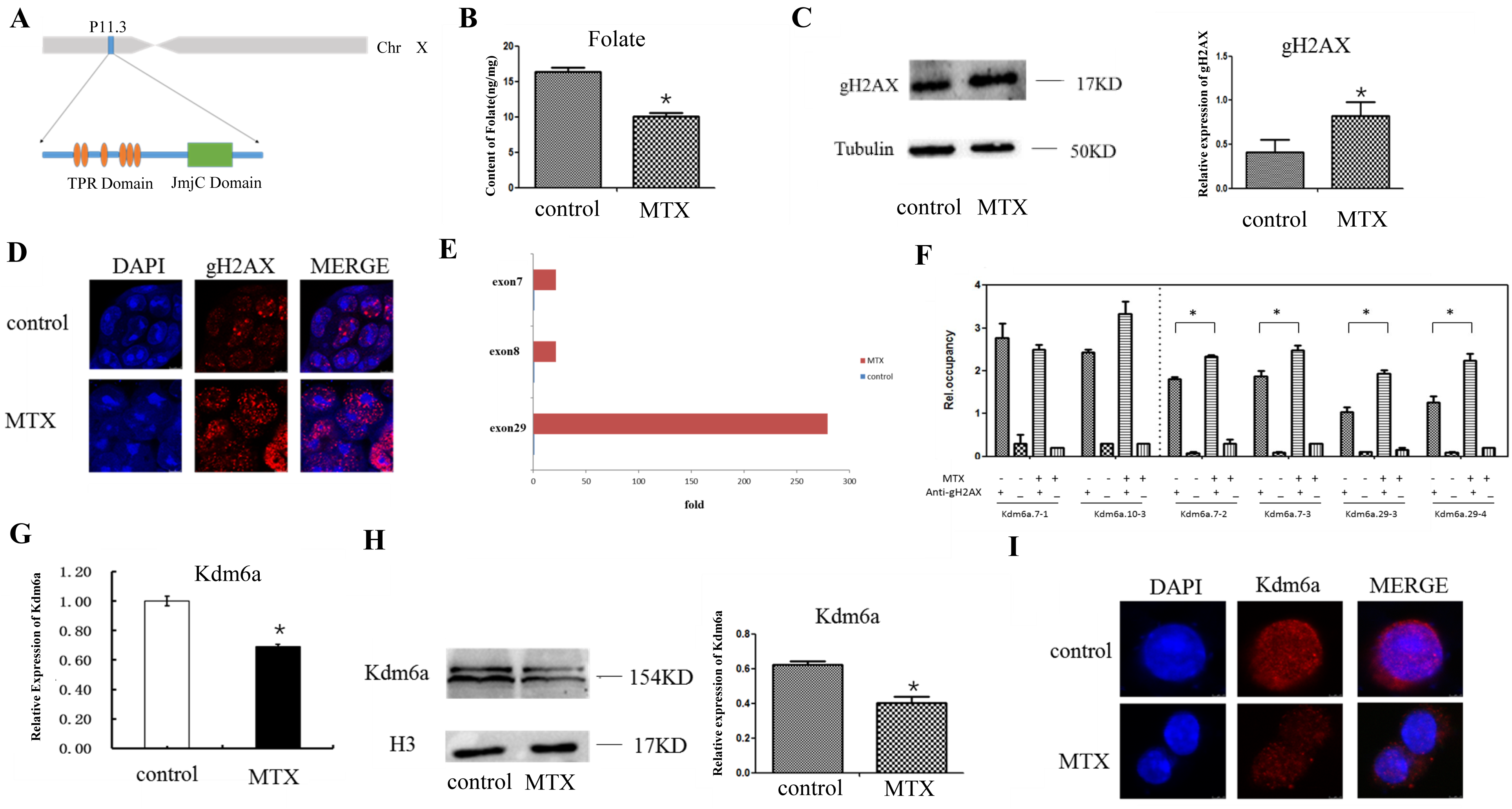
Establishment of a folate deficiency environment to study the expression of KDM6A and gH2AX. (A) Schematic diagram of the *Kdm6a* gene structure, including six tetratricopeptide repeat regions (TPR) and a transcription factor regions from jumonji family (JmjC). (B) Comparison of folate content between MTX-treated and control groups; folate was lower in the MTX group than in the normal group. (C) Western blot analysis of gH2AX protein. gH2AX was increased in the MTX group and the experiment was repeated three times. The expression was compared *p*<0.05, and the right picture was the expression comparison chart. (D) gH2AX immunofluorescence staining of the two groups. Blue indicates DAPI staining, red indicates gH2AX staining. There is stronger staining in the MTX group compared with the normal group. (E) The fracture of Kdm6a gene, blue represents the normal group and red represents the MTX-treated group, and MTX group has obvious rupture in Kdm6a, and there is almost no normal group. (F) KDM6A ChIP-qPCR map of gH2AX. The left side of the dotted line is a negative control, the difference is not statistically significant. The four groups on the right side of the dotted line are positive controls, *p*<0.05. (G) qPCR analysis of *Kdm6a*; the left side represents the normal group, the right side represents the MTX group and expression in the MTX group is lower than that in the normal group, *p*<0.05. (H) Western blot analysis of KDM6A. KDM6A expression in the MTX group was lower than that in the normal group. The experiment was repeated three times, *p* <0.05, the right picture is the expressed comparison chart. (I) Immunofluorescence staining of KDM6A. Blue indicates DAPI staining, red represents KDM6A. The fluorescence intensity was lower in the MTX group compared with the normal group.

### Reduced levels of KDM6A redistribute on the genome and participate in the regulation of NHEJ repair pathway gene expression

To investigate the reduced distribution of KDM6A on the genome, we performed KDM6A ChIP-seq. First, the signal intensity of KDM6A on gene promoter transcriptional start sites (TSSs) ± 2 kb was reproducible and correlated between the normal group and the MTX-treatment group at the genome-wide extent, Pearson coefficients (*R*)=0.6, *p*< 10^−6^ and *R*=0.63, *p*<10^−6^, respectively (Fig 2A). Next, we analyzed KDM6A enrichment in the promoter regions of NHEJ repair pathway genes. After MTX treatment, increased KDM6A binding was observed within 1 kb of the promoters of *Dclre1c, Fen1, Lig4, Mre11a, Poll, Polm, Prkdc* and *Rad50* genes, all genes in the NHEJ repair pathway (Fig 2B). To confirm the binding of Ku70, Ku80 and XRCC4 with KDM6A, we performed ChIP-qPCR and showed increased binding in the MTX group (Fig 2C). Next, we performed RT-PCR for *Ku70, Ku80* and *Xrcc4*. We found increased mRNA levels of *Ku70, Ku80*, and *Xrcc4* in the MTX group (Fig 2D). We then showed by western blotting and immunofluorescence that the nuclear levels of Ku70, Ku80 and XRCC4 proteins were increased in the MTX group compared with the control group (Fig 2E-I). The above experiments showed that in the MTX group, the expression of the NHEJ repair pathway genes, *Ku70, Ku80* and *Xrcc4*, was increased, indicating that DNA fragmentation promoted their expression, consistent with expectations. gH2AX recruits Ku70 and Ku80 to DNA breaks; therefore, we performed immunofluorescence experiments to determine gH2AX, Ku70 and Ku80 localization. We and founded that gH2AX colocalizes with Ku70 and Ku80, especially in the MTX group (Fig 2J and K), indicating that gH2AX binds to Ku70 and Ku80 at DNA breaks. The above experiments indicate that MTX induced a low folate environment in mESCs and caused DNA fragmentation. KDM6A was then enriched in the promoter regions of NHEJ repair pathway genes and increased their expression. The upregulated proteins were then recruited by gH2AX to the DNA breaks to initiate the repair process.

**Fig 2.**
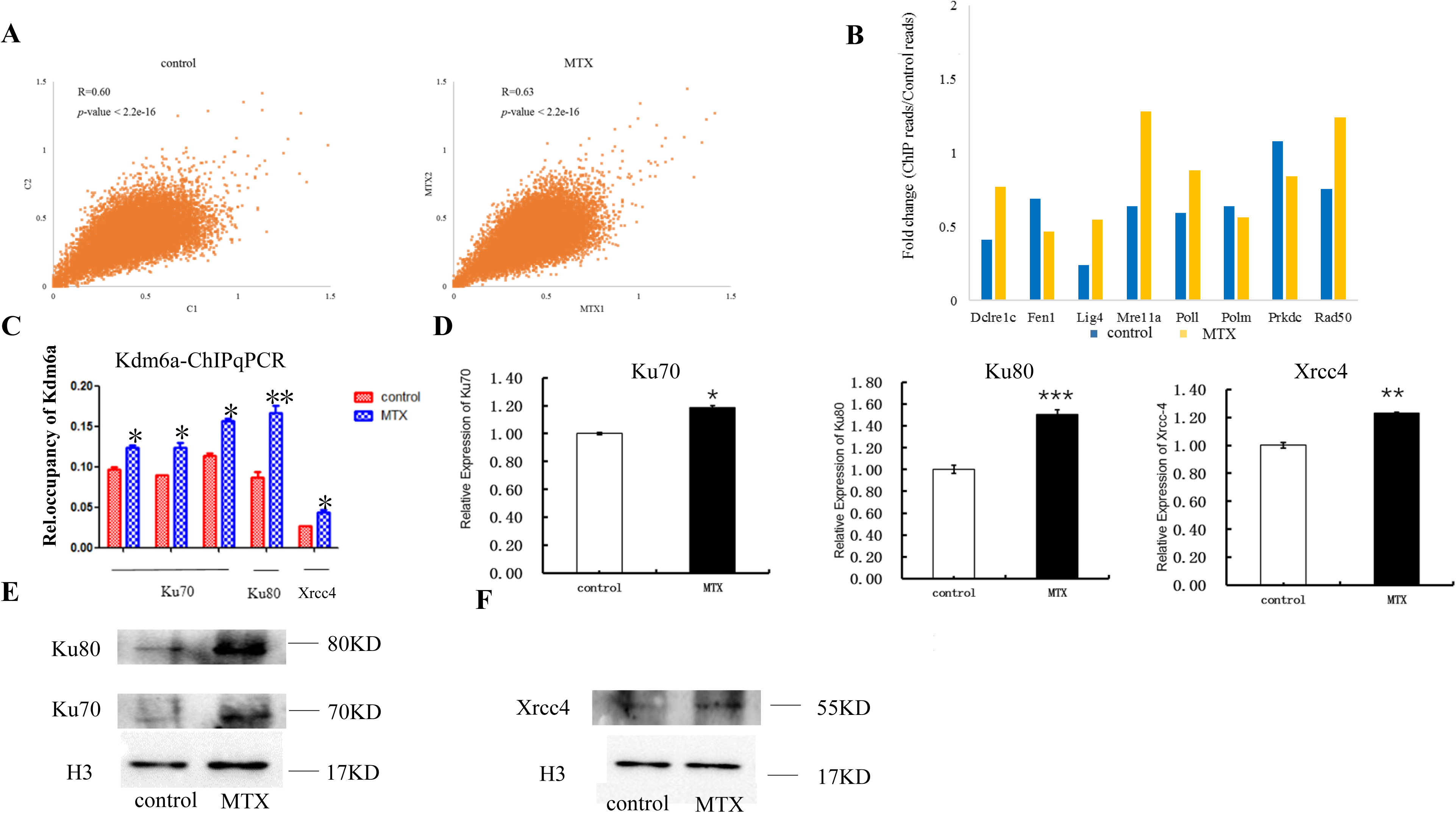
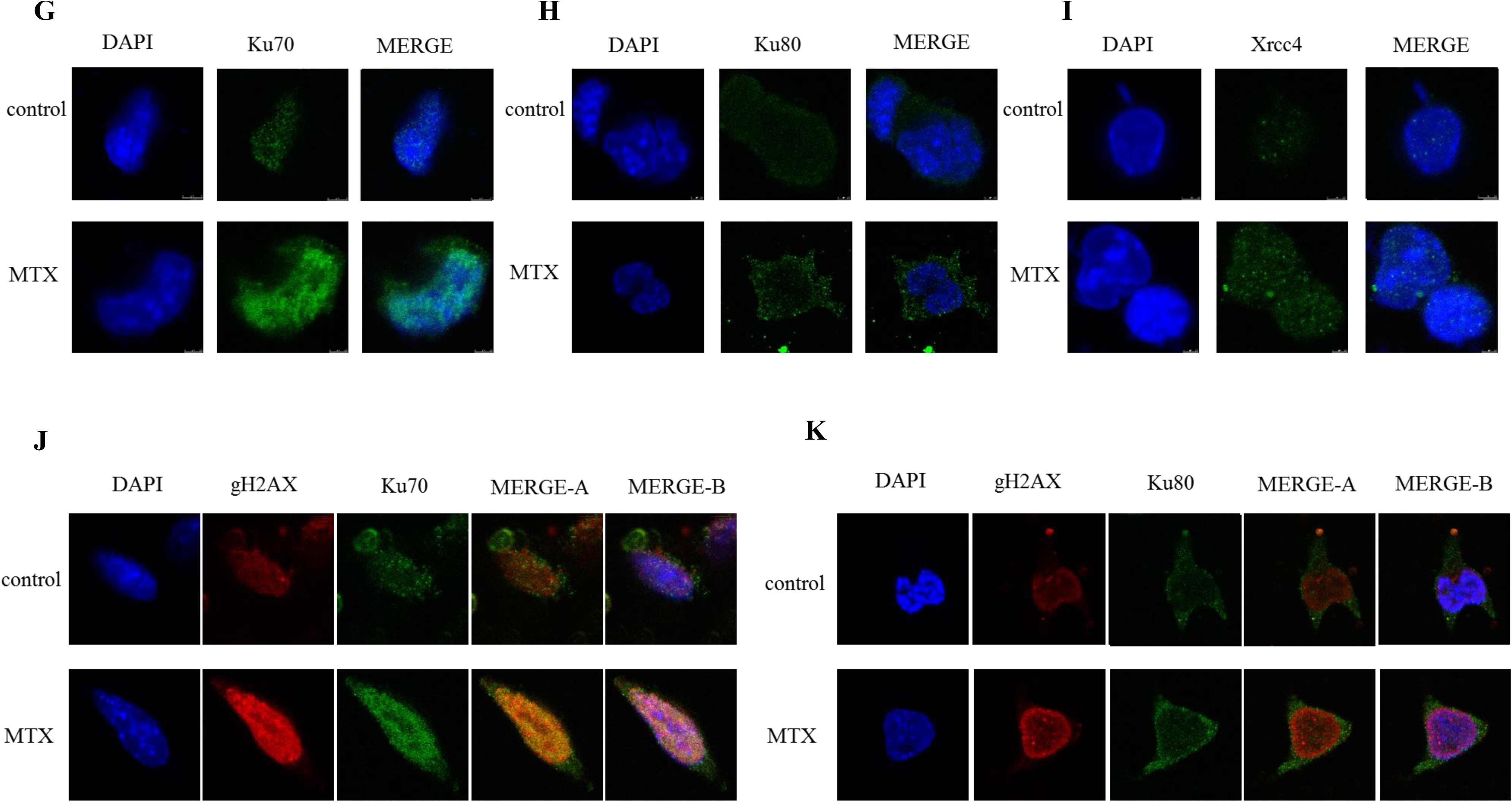
Enrichment of KDM6A on the genome and expression of corresponding proteins. (A) The signal intensity of KDM6A in genome-wide gene promoter TSSs ± 2 kb. The left side is the normal group, the right side is the MTX group, and the Pearson coefficients are *R*=0.6, *p*<10^−16^ and *R*=0.63, respectively, *p*<10^−16^. (B) Map of KDM6A binding within 1 kb of the NHEJ repair-related gene promoter regions. KDM6A binding is increased on most genes in the MTX group. (C) Ku70, Ku80 and XRCC4 ChIP-qPCR maps of KDM6A. KDM6A binding was increased on these three genes in the MTX group. Each pair of primers was tested three times, *p*<0.05. (D) RNA was extracted and qPCR was used to detect the transcription of *Ku70, Ku80* and *Xrcc4*. The transcription of these three genes was increased in the MTX group. (E) Western blot of Ku70 and Ku80. The expression of both proteins was increased in the MTX group. (F) Western blotting showed increased expression of XRCC4 in the MTX group. (G-I) Immunofluorescence of Ku70 (G), Ku80 (H) and XRCC4 (I) in mESCs. Blue represents DAPI, and green represents Ku70, Ku80 and XRCC4, respectively. The fluorescence signal was increased in the MTX group. (J-K) Immunoconfocal images of Ku70 (J) and Ku80 (K) in mESCs cells, where blue represents DAPI, red represents gH2AX, and green represents Ku70 and Ku80. MERGE-A represents a Ku (70/80) fusion map with gH2AX, MERGE-B is a fusion map of Ku (70/80) with gH2AX and DAPI. Both proteins were increased in both the MTX group.

### KDM6A regulation of NHEJ repair pathway activity is dependent on p53

As a key factor in the DNA repair process, p53 plays an essential role in activation of the DNA break repair pathway. Here, we explore further its role in the activation of the NHEJ pathway. p53 binds to KDM6A in *Drosophila* to activate the NHEJ pathway. To investigate whether this also occurs in mESCs, we used p53siRNA (50 nM) to knock down the expression of p53 (Fig 3A) in cultured cells. Forty-eight hours after electroporation of p53siRNA, MTX (0.12 μM) was added to cells. Proteins were then isolated for p53 Co-IP experiments. We did not identify KDM6A in the group without any MTX treatment. However, in the MTX-treated group, the level of KDM6A was increased (Fig 3B), indicating that MTX enhanced binding between p53 and KDM6A after DNA fragmentation. Next, we kept cells with no behavior and p53siRNA electroporation. and added MTX to both cells respectively. KDM6A was almost absent in cells transfected with p53siRNA (Fig 3C), indicating that the binding of KDM6A to p53 was decreased and that the binding of p53 to KDM6A is dependent on the amount of p53. We also performed immunofluorescence staining for p53 and KDM6A. In the MTX group, the level of KDM6A and p53 binding was enhanced, while in no behavior and p53siRNA electortransfered the nuclear binding of these two proteins was weak (Fig 3D). This phenomenon is consistent with the above experimental results. We then explored further the effect of KDM6A on the binding of Ku70, Ku80 and XRCC4 after knocking down p53. ChIP for KDM6A showed that KDM6A reduced the binding among Ku70, Ku80 and XRCC4 in the p53 group compared with the control group (Fig 3E). This result indicates that p53 has an effect on the NHEJ repair pathway. When p53 binds to KDM6A to form a complex, it promotes the expression of NHEJ repair pathway genes. After knockdown of p53, the binding of KDM6A to NHEJ repair proteins is weakened and inhibits their expression.

**Fig 3.**
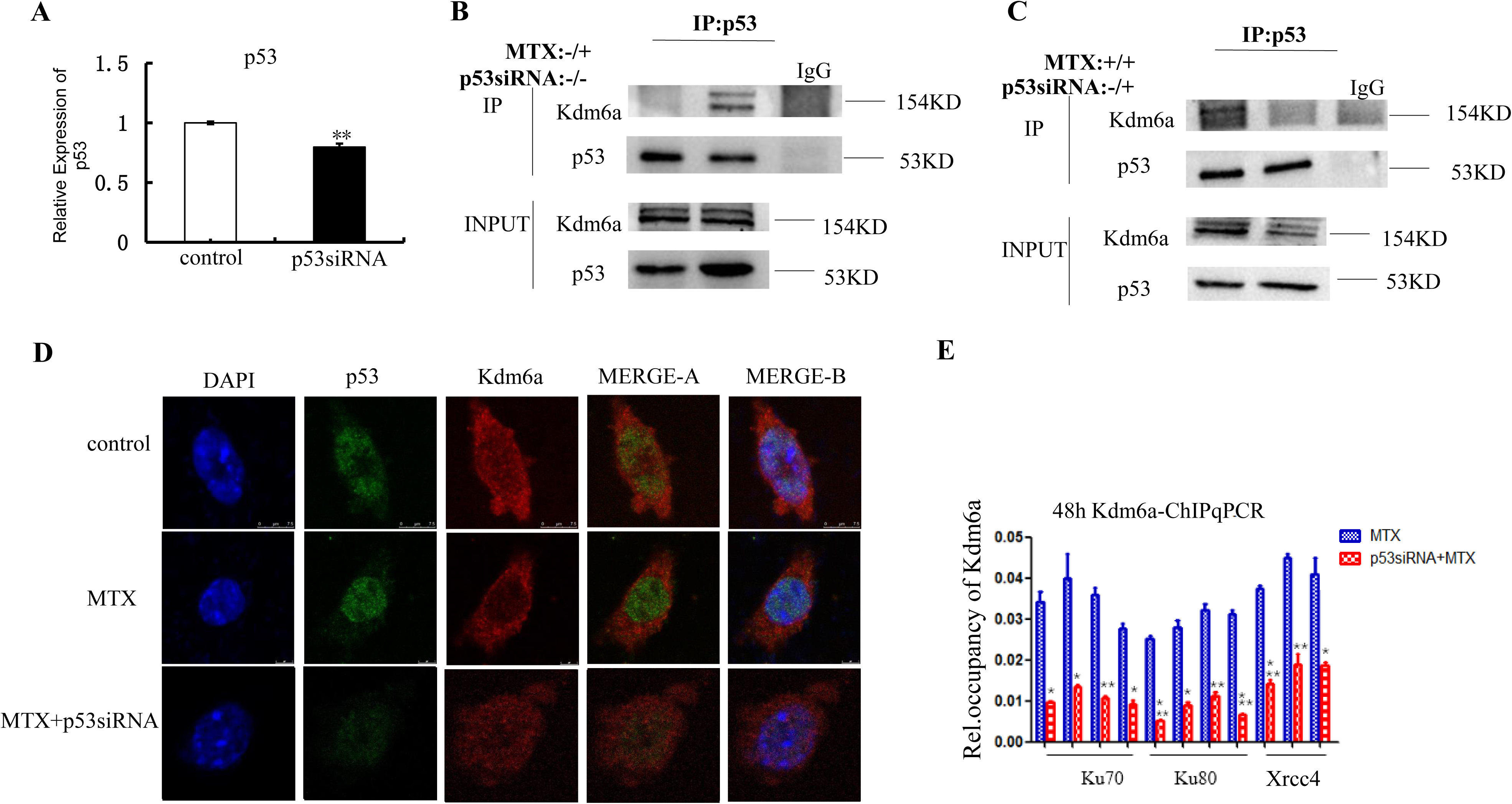
Interaction of p53 and KDM6A in DSBs. (A) qPCR was performed 48 h after electroporation of mESCs with p53siRNA, and the transcription level of p53 was decreased. (B) p53 and KDM6A co-immunoprecipitation (Co-IP) using a p53 antibody. p53 and KDM6A were weakly bound in the normal group (left) and more strongly bound in the MTX group (middle). IgG was used as a negative control (right). (C) Cells were added MTX and divided p53siRNA transfected (experimental group) and no transfected (blank group). After 48 hours, proteins were extracted for Co-IP, and the blank group p53 and Kdm6a were distinct (left), experimental group p53 and Kdm6a binding wasn’t obvious (middle), IgG was used as a negative control (right). (D) Cells were divided three groups, no behavior group (control), added MTX and transfected p53siRNA group (MTX+p53siRNA), and only added MTX group (MTX). Three groups were performed immunoconfocal of p53 and Kdm6a, in the MTX group, the nuclear trajectories of p53 and Kdm6a are obvious (middle), while in control (upper) and MTX+p53siRNA group (bottom) p53 and Kdm6a bindings were not obvious. MERGE-A represents the fusion map of p53 and Kdm6a, and MERGE-B represents the fusion map of p53 and Kdm6a and DAPI. (E) MTX was added to the transfected p53siRNA 24 h experimental group and the untransfected control group, respectively. After 24 hours, a Kdm6a-ChIP experiment was performed, followed by qPCR analysis of Ku70, Ku80 and Xrcc4.The binding of Kdm6a with Ku70, Ku80 and Xrcc4 in the experimental group is reduced.

### Decreased *Kdm6a* expression associates with increased *Ku80* expression in NTDs

We verified that the expression level of *Kdm6a* in mESCs was decreased in an MTX-induced low-folate environment, and that the expression level of Ku80 was increased. Changes in protein levels in NTD samples and in the mouse NTD model were examined. Development of the mouse nervous system is complete by E10.5; therefore, we used this time point to select embryos. Normal embryos were intact and morphologically smooth, while NTD embryos showed a bulge in the brain (Fig 4A). We performed immunohistochemical analysis of KDM6A in normal and NTD embryos. KDM6A levels were decreased in embryonic brain and spinal tissues in NTD samples (Fig 4B).

**Fig 4.**
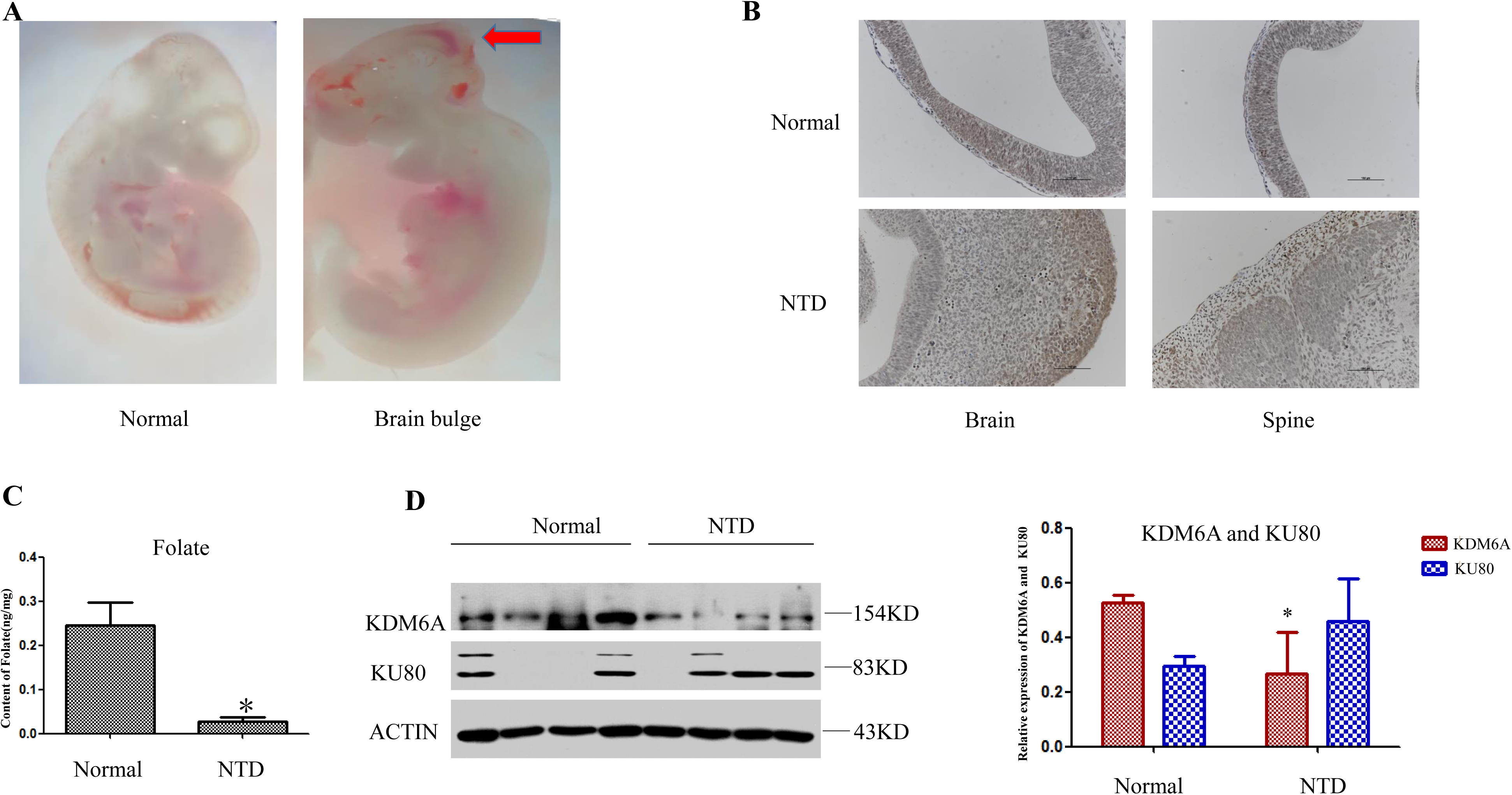
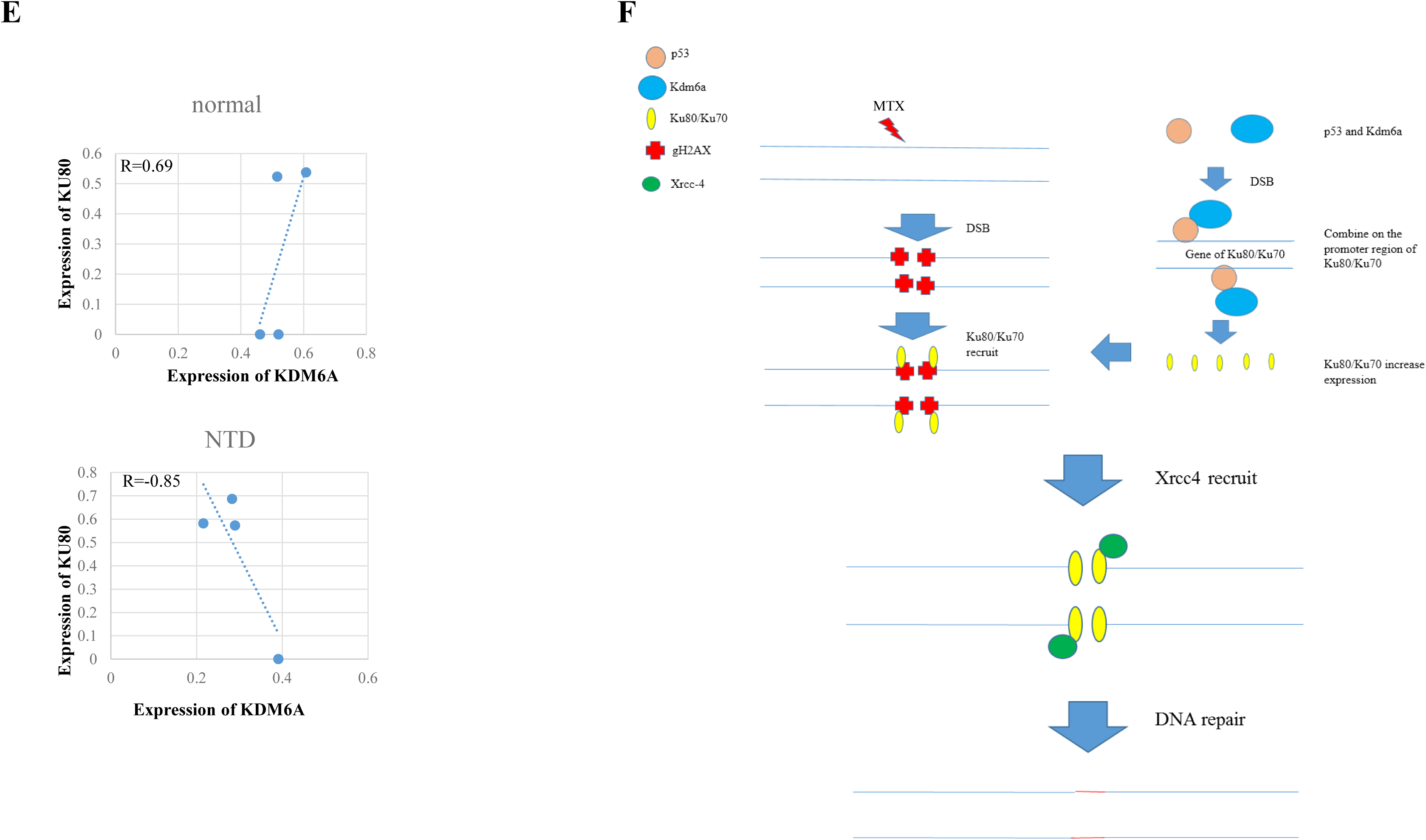
Expression levels of KDM6A and KU80 in NTDs. (A) Mouse embryos 10.5 days during pregnancy were observed under the microscope after RA induction. The left image shows a normal mouse embryo, and the right image shows an NTD mouse embryo. It was marked with an arrow, and there was a bulge of embryonic brain. (B) KDM6A immunohistochemistry in mouse embryos with RA-induced at E10.5, and in selected brain tissues (left) and spinal tissue (right). Normal tissue staining was intense (top), and weak in NTDs (bottom). (C) Determination of folate content in normal human specimens and NTDs specimens. The folate content in NTD tissues was low, *P* < 0.05. (D) Western blot analysis of normal human tissues and NTD tissues was performed to analyze the levels of KDM6A and KU80. The level of KDM6A in NTD samples was lower than that in normal tissues, *p*<0.05. KU80 expression was increased in NTD specimens. The right panel is an expression of the two proteins. (E) Schematic representation of the correlation of KU80 and KDM6A expression. In normal samples, the expression of the two proteins was positively correlated, *R*= 0.69 (top). In NTD specimens, the expression of the two proteins was negatively correlated, *R*= −0.85 (bottom). (F) MTX induces DSBs, and the NHEJ repair pathway is initiated to repair the broken DNA.

To explore the mechanism of KDM6A action during the development of human NTDs, we hypothesized that early intake of folate in early stages of embryonic development may lead to disordered regulation of KDM6A, resulting in abnormal embryo development. Therefore, we selected brain tissues from four normal and four NTD embryos for analysis. Embryo information is presented in Supplementary Table S3. The folate level of the NTD group was 0.035 ± 0.024 ng/10^6^, which was significantly lower than 0.272 ± 0.119 ng/10^6^ of the normal group (Fig 4C). Western blot analysis of human samples demonstrated KDM6A was in the NTD group was half that compared with the normal group, while the content of KU80 was higher in the NTD group than in the normal group (Fig 4D). Correlation analysis showed a positive correlation between the two in normal tissues (Pearson value 0.69); and there was a negative correlation in NTD tissues (Pearson value −0.85) (Fig 4E). The above results indicate the possibility that KDM6A levels are decreased and KU80 levels are increased in folate deficient conditions, which perturbs normal nervous system development of the embryo, leading to the occurrence of NTDs. Together, these data show that MTX treatment creates a low-folate environment resulting in DSBs, which leads to KDM6A binding to p53 to form a complex that activates the NHEJ pathway to repair the DSBs (Fig 4F).

## Discussion

The integrity and stability of the structure of DNA is crucial to cell function and survival. Various endogenous and exogenous factors can cause different forms of DNA damage. Various physical, chemical and metabolic toxins may cause structural damage to chromosomes and mitochondrial DNA, including single and double strand breaks and chemical crosslinks between DNA and protein molecules. Among them, DSBs are the most serious form of damage (Lengauer et al., 1998; Pastink and Eeken, 2001; Stucki et al., 2005). Unrepaired DSBs lead to further genomic damage including chromosome translocation, degradation and amplification. These can affect the copy number of genes and dysregulate gene expression, leading to diseases, such as tumors, immunodeficiency diseases, radiation sensitivity disorders and neurodegeneration (Larsen and Stucki, 2016; PaigenK and PetkovP, 2010; JacksonSP and BartekJ, 2009; Branzei, 2010).

Development of the vertebrate central nervous system is a rigorous, orderly and complex process. The nervous system of mammals is very sensitive to DSBs during embryonic development. When DSBs occur, cells need to act rapidly to repair them. Our present study reveals a novel role for KDM6A in maintaining genomic stability. We show in mESCs that KDM6A physically interacts with p53 under folate-deficient conditions, but that there is no interaction between them under normal folate levels. These findings indicate that p53 may be involved in the activation of core NHEJ repair factors that are demethylated by KDM6A depending on the cellular folate level.

p53 plays an important role in DNA damage-induced G2/M checkpoint activation and in inhibiting entry into mitosis (Fernandez-Capetillo et al, 2002; Yang et al.2018). In our previous report, we established the range of MTX doses that induces DNA breaks in mESCs. A remarkable decrease in S phase cell numbers (about 30%) was visible with 0.12 μM MTX, indicating transition to the G1/S checkpoint was delayed. The proportion of cells in G2/M was remarkably declined compared with the normal group. Also no obvious apoptosis was detected at 0.12 μM MTX, excluding the likelihood that the extensive fragmentation of nuclear DNA was the results of apoptosis. However, the precise mechanism of p53 action is still not clear and needs further study.

Animal experiments have shown that defects in DNA repair genes can affect the survival of embryos and cause birth defects. Murai et al found that mice with a defective NER pathway had severe neurodevelopmental abnormalities, such as decreased cerebellum volume, damaged Purkinje cell dendrites, slowed proliferation of granular layer neurons and increased apoptosis (Murai et al., 2001). In our study, we found a compensatory effect of DNA repair via the NHEJ pathway after folate deficiency *in vitro* and *in vivo*. It is noteworthy that we characterized brain tissue of NTD-affected fetuses from gestational weeks 20–36 instead of from the first trimester, which is considered to be the critical period for neurulation. Mouse nervous system development is complete at E10.5 (Copp and Greene, 2010; Massarwa and Niswander, 2014); therefore, we examined neural tissue from the NTD mouse model at this time-point. Immunohistochemistry showed KDM6A levels were decreased in the brains of NTD mouse embryos compared with normal mouse embryos, indicating the down-regulation of KDM6A during neural tube closure. Nevertheless, the findings of this study reveal a previously uncharacterized role of KDM6A in stabilizing the genome under folate deficient conditions during development of the nervous system.

In summary, DNA breaks trigger KDM6A to interact with p53, which modulates expression of the Ku heterodimer (Ku70/80) and XRCC4, to ultimately promote the DNA repair response after folate deficiency. This study advances our knowledge of KDM6A in the maintenance of genomic integrity and provides a new understanding of NHEJ repair pathway regulation in the etiology of NTDs.

## Materials and Methods

### Cell Culture and Folate Treatment

SV129 mESCs were obtained from the Stem Cell Bank, Chinese Academy of Sciences (Shanghai, China). After seeding onto culture dishes coated with 0.2% gelatin (Sigma-Aldrich, St. Louis, MO USA), cells were incubated in medium (Invitrogen, Carlsbad, USA) supplemented with 0.1 mM non-essential amino acids (Invitrogen), 0.1 mM glutamate (Invitrogen), 0.1 mM β-mercaptoethanol (Life Technologies), 1000 U/ml leukemia inhibitory factor (Millipore, USA), and 15% fetal bovine serum (Invitrogen). Cells were maintained at 37°C in a humidified atmosphere with 5% CO2 and passaged every 3 days. The medium was changed daily. When cells reached 75% confluency, 0.12 μM MTX (Sigma-Aldrich) was added to the culture flask. Cells were collected 24 hours later.

### Immunofluorescence

Medium was removed and cells were washed twice with PBS. Cells were fixed in 200 μl of chilled 4% paraformaldehyde (Solarbio, Beijing,China) at room temperature for 10 min. Cells were then washed three times with 500 μl PBS for 5 min each time. Cells were permeabilized for 10 min in 200 μl 0.5% Triton X-100 in PBS (Solarbio) and then rinsed with PBS at room temperature for 5 min. Cells were then blocked with 500 μl immunofluorescence blocking solution (1×) per well at room temperature for 60 min. The blocking solution was removed and cells were then incubated with 250 μl of the primary antibody dilution at 4°C overnight. Cells were washed in 500 μl PBS for 5 min, three times. Cells in each well were then incubated with 250 μl secondary antibody dilution for 1 h at room temperature and protected from light. The secondary antibody was removed and cells were washed three times with 500 μl PBS for 5 minutes each time. Cells were then stained with DAPI, stored in the dark, and observed under a fluorescence microscope (Zeiss LSM710).

### Western Blot Analysis

Proteins were extracted from SV129 mESCs and tissues using a protein extraction kit (C500009; Sangon Biotech, Shanghai, China), following the manufacturer’s protocol. Cells or tissues were washed twice with PBS, mixed with extraction reagent and left to stand for 10 minutes. Samples were then centrifuged at 3500 rpm for 5 minutes. The supernatant was discarded and 200 μl of lysate was added. After incubation on ice for 20 minutes samples were centrifuged at 12000 rpm for 10 minutes at 4°C. The supernatant containing nuclear proteins was stored at −80°C. For western blotting, nuclear proteins were subjected to 10% SDS–polyacrylamide gel electrophoresis and transferred to a polyvinylidene difluoride membrane. The membrane was then blocked with 5% non-fat milk in PBS and 0.1% Tween 20 (PBS-T) for 1 h, followed by incubation with the primary antibody at 4°C overnight and then with the secondary antibody for 1 h. Detection was performed using Amersham™ ECL™ Prime Western Blotting Detection Reagent (GE Healthcare) and a SuperSignal West Femto Trial Kit (Thermo Scientific, USA).

### Co-immunoprecipitation

Co-immunoprecipitation (Co-IP) experiments were performed using a Crosslink IP kit (Thermo Fisher) according to the manufacturer’s instructions. Five micrograms of antibody were bound to protein A/G plus agarose for 45 minutes at room temperature with mixing on a rotary mixer. An IgG negative control was also prepared. Bound antibody was then reacted with 450 μM disuccinidyl on a rotary mixer at room temperature and the aminosuccinate then crosslinked for 50 minutes. At the same time, pre-clearing of the nuclear protein using control agarose resting on a rotator was performed at 4 °C for 50 min. The pre-eluted nucleoprotein was then incubated with the antibody-crosslinked resin overnight at 4°C to complete antigen immunoprecipitation. Antigen elution was completed the next day and the bound protein was detected by western blot analysis.

### Chromatin Immunoprecipitation (ChIP), ChIP-qPCR, and ChIP-seq

Enzymatic Chromatin IP Kit (Cell Signaling Technology, MA, USA) was used for ChIP assays according to the manufacturer’s instructions. Chromatin was obtained from about 2×10^7^ SV129 mESCs and crosslinked with formaldehyde. Immunoprecipitation was then performed overnight at 4°C with a KDM6A antibody. The immunoprecipitated DNA was analyzed by in-depth genome-wide DNA sequencing, performed by Capital Bio Technology (Beijing, China). Raw sequencing data were examined using the Illumina Analytical Pipeline, aligned with the *Mus musculus* reference genome (UCSC, mm10) using Bowtie2, and further analyzed by MACS (model-based ChIP-Seq analysis; https://github.com / taoliu / MACS). DNA-protein complexes were analyzed by qPCR with specific primers to amplify multiple regions of the gene. Several pairs of primers were designed. qPCR was performed using SYBR SuperMix (TransGen Biotech, Beijing, China) according to the manufacturer’s instructions (The primers used are listed in Supplementary Table S1.). Amplification data acquisition and analysis were performed using the 7500 Fast Real-Time PCR System (Applied Biosystems). The percentage of DNA captured by ChIP (input percentage) was calculated as follows: Percent input = 2% × 2(^C[T] 2% input sample – C[T] IP sample^), where C[T] = Ct = threshold cycle of the PCR reaction. Three independent ChIP experiments were performed for each analysis.

### RNA Extraction, Reverse Transcription, and Real-time RT-PCR Analysis

Total RNA was isolated from 0.5×10^6^ cells using TRIzol reagent (Invitrogen, Carlsbad, CA, USA). Total RNA from each group was converted to cDNA using TransScript First-Strand cDNA Synthesis SuperMix (TransGen Biotech, Beijing, China). Real-time PCR to quantify relative mRNA levels in different cell groups was performed using UltraSYBR Mixture (CW Biotech, Beijing, China) and the QuantStudioTM 7 Flex Real-Time PCR System (ABI, CA, USA). Primers were synthesized by CW Biotech (Beijing, China). The reaction composition was as follows: forward and reverse primers (0.5 μmol, 10 μmol, respectively). (The primers used in this experiment are listed in Supplementary Table S2.), UltraSYBR mixture (12.5 μl), cDNA (1 μl) and ddH_2_O (10.5 μl). Amplification parameters were as follows: 3 minutes of initial denaturation at 95°C, 45 amplification cycles of denaturation at 95°C for 15 seconds, annealing at 58°C for 20 seconds, extension at 72°C for 30 seconds, and then one cycle at 72°C for 10 minutes. Gene expression was normalized to the expression of *Gapdh*. The relative levels of mRNA transcripts were calculated using the classical ΔΔCt method.

### Nuclear Transfection

Nuclear transfection of SV129 mESCs was performed using an Amaxa^®^ Nucleofector II Device (Lonza, Sweden) according to the manufacturer’s instructions. First, cells were collected and centrifuged at 1000 rpm for 5 minutes at 4°C. The supernatant was discarded and 90 μl of electroporation solution and p53 siRNA (#6231; CST) added to the tube and mixed. and transferred together, running on the Amaxa Nucleofector program Performed on an electric rotator under A-023 item of the Amaxa Nucleofector. After completion, mESCs were transferred to preheated medium for cultivation.

### Retinoic Acid Treatment of Mouse Embryos

KM mice were obtained from Beijing Vital River Laboratory Animal Technology Co. Ltd. Mature male and female KM mice were mated overnight and the presence of vaginal plugs detected the next morning. The presence of a vaginal plug indicated a time point of 10.5 days during embryogenesis (E10.5). Retinoic acid (RA) (Sigma) was dissolved in olive oil at a concentration of 20 mg/kg and injected intraperitoneally on E7.5. Control mice were given 20 mg/kg olive oil intraperitoneally. At E10.5, pregnant mice were euthanized by cervical dislocation. Embryo phenotypes were observed under a microscope, and brain and spinal cord tissues of normal and spina bifida phenotype embryos were dissected as previously reported (Li et al. People, 2015; Yu et al., 2017). All animal experiments were approved by the Animal Ethics Committee of the Capital Pediatric Society.

### Immunohistochemistry

Paraffin sections were placed in an oven at 67°C, baked for 2 hours, and rinsed three times with PBS for 3 minutes each. Antigen retrieval was then performed by boiling samples in citrate buffer in a microwave. Slides were then rinsed twice with distilled water and then three times with PBS. One drop of 3% H2O2 was then placed on each section, incubated for 10 minutes at room temperature, and slides were rinsed three times with PBS for 10 min each time. One drop of KDM6A antibody dilution was then placed on each section, and incubated for 2 hours at room temperature. After rinsing three times with PBS for 5 min each time, 1 drop of polymer enhancer was added to each section and incubated for 20 minutes at room temperature. Slides were then rinsed three times with PBS for 3 min each time and 1 drop of the enzyme-labeled anti-mouse/rabbit polymer added to each section and incubated for 30 minutes at room temperature. After rinsing three times with PBS for 5 min each time, 1 drop of freshly prepared DAB solution (diaminobenzidine) was added to each section and staining was monitored under a microscope for 5 minutes. Next, hematoxylin counterstaining, 0.1% HCl differentiation, tap water washing, bluening, sectioning and dehydration drying by gradient alcohol, xylene transparent, neutral gum sealing were performed, drying and then observed.

### Determination of Folate Concentration

SV129 cells were harvested and folate concentrations were determined using a competitive receptor binding immunoassay (A14208, Chemiluminescent Immunoenzyme Assay; Beckman Coulter, Fullerton, USA) and the Access 2 Immunoassay system (Beckman Coulter) according to the manufacturer’s instructions. Cells were collected in 1 ml of TRIS buffer, sonicated for nine cycles and centrifuged at 10,000 rpm for 3 minutes at 4°C. The supernatant was collected for folate measurement.

### Antibodies

The primary antibodies used were purchased from the following sources: anti-KDM6A (#ab36938, 1:500; Abcam); anti-gH2AX (#D2893, 1:1000; Abcam); anti-KU80 (#ab119935, 1:1000; Abcam); anti-KU70 (sc-17789;1:500,Santa Cruz), anti-XRCC4 (#sc-271087, 1:500; Santa Cruz); anti-p53 (#2524, 1:1000; CST); anti-histone H3 (#ab21054, 1:5000;Abcam); anti-alpha-tubulin (#ab7291, 1:100; Abcam); anti-beta-actin (#TA-09, ZSGB-BIO; 1:1000).

### Statistical Analysis

All data are expressed as the mean ± s.d. using SPSS 22.0 software. A two-tailed t-test was performed to analyze the difference between two groups, with statistical difference set at *P* < 0.05.

## Subjects

All clinical samples were from patients from Shanxi Province, China. The families of all patients gave informed consent. Diagnosis was performed by clinical ultrasound. The ethics committee of the Capital Pediatrics Society approved this study.

## Acknowledgements

We are grateful for the help provided by Weifang Medical College and the Capital Institute of Pediatrics. We thank Jeremy Allen, PhD, from Liwen Bianji, Edanz Group China (www.liwenbianji.cn/ac), for editing the English text of a draft of this manuscript.

## Funding

This study was supported by CAMS Initiative for Innovative Medicine (2016-I2M-1-008), Beijing municipal program of medical research (Grant No. 2016-04), the National Natural Science Foundation of China, Beijing, China (81771584), and the Pediatric Medical Coordinated Development Center of Beijing Municipal Administration of Hospitals (XTZD20180402).

## Competing interests

The authors declare that they have no conflict of interest.

